# A Novel *Ex Vivo* Peritoneal Model to Investigate Mechanisms of Peritoneal Metastasis in Gastric Adenocarcinoma

**DOI:** 10.1101/2021.11.15.468687

**Authors:** Deanna Ng, Aiman Ali, Kiera Lee, Denise Eymael, Kento Abe, Shelly Luu, Karineh Kazazian, Savtaj Brar, James Conner, Marco Magalhaes, Carol J Swallow

**Affiliations:** Institute of Medical Science, University of Toronto, Toronto, Canada; Lunenfeld-Tanenbaum Research Institute, Sinai Health System, Toronto, Canada; Faculty of Dentistry, University of Toronto, Toronto, Canada; Department of Surgical Oncology, Princess Margaret Cancer Centre/Mount Sinai Hospital, Toronto, and Department of Surgery, University of Toronto, Toronto, ON, Canada; Department of Pathology, Mount Sinai Hopsital

**Author notes:** Please address correspondence to: Carol J. Swallow MD PhD, Mount Sinai Hospital, 600 University Avenue #1225, Toronto, ON Canada M5G 1X5.

**Keywords:** *Ex Vivo* model, Peritoneal Metastasis, Gastric Cancer

## Abstract

Peritoneal metastases (PM) portend limited survival in patients with Gastric Adenocarcinoma (GCa), and strategies to prevent and/or more effectively treat PM are needed. Existing models are limited in recapitulating key elements of the peritoneal metastatic cascade. To explore the underlying cellular and molecular mechanisms of PM, we have developed an *ex vivo* human peritoneal explant model. Fresh peritoneal tissue samples were obtained from patients undergoing abdominal surgery and suspended, mesothelial layer down but without direct contact, above a monolayer of red-fluorescent stained AGS human GCa cells for 24hrs, then washed and cultured for a further 3 days. Implantation and invasion of GCa cells within the explant were examined using real-time confocal fluorescence microscopy. Superficial implantation of AGS GCa cells within the mesothelial surface was readily detected, and colonies expanded over 3 days. To investigate the sensitivity of the model to altered GCa cellular implantation, we stably transfected AGS cells with E-Cadherin, restoring the E-Cadherin that they otherwise lack. This markedly suppressed implantation and invasion of AGS cells into the submesothelial mesenchymal layer. Here we show that this *ex vivo* human peritoneal explant model is responsive to manipulation of genetic factors that regulate peritoneal implantation and invasion by GCa cells, with reproducible results.

## INTRODUCTION

The peritoneal cavity is a potential space that is enveloped in a thin layer of specialized tissue known as the peritoneum [1, 2]. The peritoneal surface can be the target of a variety of inflammatory, infectious, and neoplastic processes, for instance peritonitis associated with perforated appendicitis, tuberculosis and peritoneal carcinomatosis, respectively [3, 4]. The predilection of certain types of primary cancer to spread within the peritoneum has long been the subject of scientific inquiry [5]. The peritoneal niche is considered a “soil” within which certain epithelial malignancies characteristically “seed”. Prominent examples include ovarian, gall bladder and gastric adenocarcinomas [6, 7]. The cellular and molecular features that enable this specialized interaction remain largely undefined.

In the case of gastric adenocarcinoma (i.e., stomach cancer) metastasis to the peritoneum remains a common and challenging problem. In North America and Europe, approximately 35% of patients who are newly diagnosed with gastric cancer already have peritoneal metastasis, and the median survival in such patients is ^~^4 months [8–10]. Of those who appear to and undergo curative intent resection, about one quarter develop peritoneal recurrence, with a median survival of 6 months even when treated with aggressive palliative chemotherapy [11, 12]. Peritoneal metastases from gastric adenocarcinoma cause ascites, ureteric and bowel obstruction, and intractable symptoms that are difficult to manage effectively [13, 14]. Peritoneal carcinomatosis degrades both quantity and quality of life with gastric cancer. Identification of the mechanisms and distinct molecular profile that predisposes to peritoneal spread will lead to discovery of rational targets for the prevention and/or treatment of this stubborn metastatic pattern.

However, the elucidation and validation of the mechanism of peritoneal metastasis have proven difficult to study. Peritoneal dissemination of cancer is a complex and dynamic process that, like the hematogenous metastatic cascade, can be conceptualized as comprising multiple steps. The peritoneal metastatic cascade has 5 major steps: 1) detachment of cancer cells from the primary tumour, 2) survival in the microenvironment of the peritoneal cavity, 3) attachment of free tumour cells to peritoneal mesothelial cells, 4) invasion into the subperitoneal space and 5) tumour growth [15–17]. As shown, the peritoneal metastatic cascade notably differs from the hematogenous and lymphatic cascades, both of which entail directional, intra- and extra-vasation through vessel walls (Fig. 1A) [18–20]. The study of established gastric cancer peritoneal metastasis, such as that shown in Fig. 1B, can be hampered by paucicellularity and surrounding fibrosis. In addition, the molecular features that enabled steps 1 through 4 of the peritoneal metastatic cascade may no longer be relevant/ represented.

**Figure 1.**
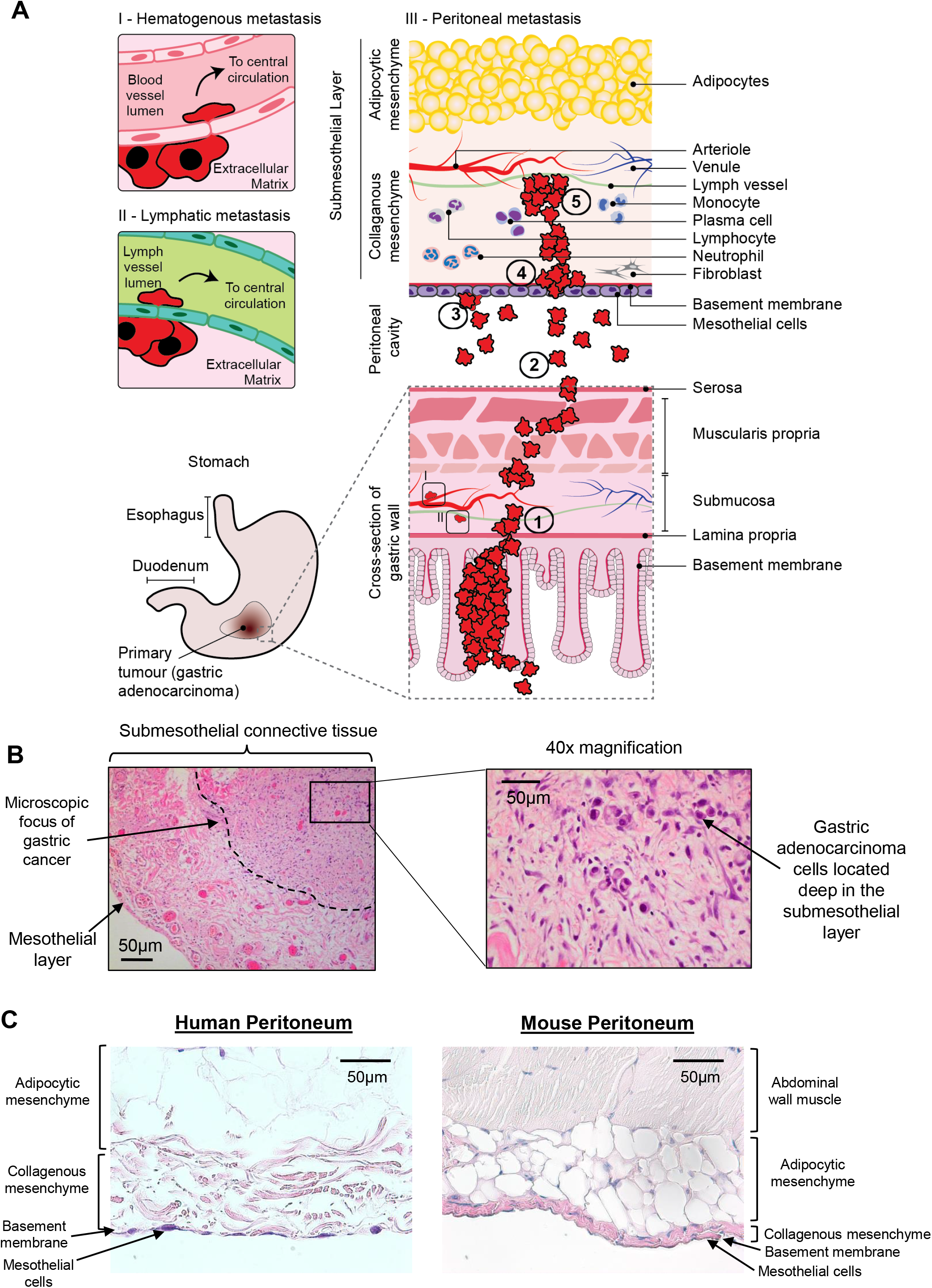
Human peritoneum represents a distinct niche for dissemination of gastric adenocarcinoma. **A)** Cartoon depicting routes of metastatic spread from the primary tumour in gastric adenocarcinoma. The hematogenous (I) and lymphatic (II) metastatic cascades involve directional intra- and extravasation of cancer cells in and out of vessels. This is not the case for the peritoneal metastatic cascade, which comprises 5 major steps: 1) detachment of cancer cells from the primary tumour at the serosal surface of the stomach; 2) survival and movement within the microenvironment of the peritoneal cavity; 3) attachment of free tumour cells to peritoneal mesothelial cells; 4) invasion into the subperitoneal layer and 5) proliferation of cancer cells within the subperitoneal mesenchyme. **B)** Metastatic deposit within the peritoneum. Cross sectional image taken perpendicular to mesothelial surface showing established gastric adenocarcinoma peritoneal metastasis deep in submesothelial layer, with surrounding fibrotic response (Dashed line depicts periphery of peritoneal metastasis). **C)** Representative images of formalin-fixed paraffin-embedded (FFPE) sections of human (left) and mouse (right) peritoneum stained using H&E (×10) illustrating significant differences in histoarchitecture, including a thinner layer of collagenous mesenchyme in the mouse.

Our current understanding of the mechanisms of peritoneal carcinomatosis in gastric adenocarcinoma has been formulated based on animal models or artificial *in vitro* cellular representations of the human peritoneum [21–23]. Most in vivo animal models do not recapitulate the entire metastatic cascade, and have the disadvantage that they employ immunodeficient animals [24–26]. Furthermore, there has been little reproducibility between studies. The marked physical differences between mouse peritoneum and human peritoneum are clear upon routine histologic assessment (Fig. 1C). The main limitation of existing *in vitro* models is that mesothelial cells are cultured in a monolayer, and this does not accurately model the behaviours of cancer cells moving within a complex 3D environment such as the peritoneal cavity [27–29].

We aimed to create a more relevant depiction of the human peritoneal cavity, by allowing gastric cancer cells to interact with explanted human peritoneum *ex vivo*. Having demonstrated the viability, specificity and reproducibility of the model in measuring implantation and invasion of human gastric cancer cells into human peritoneum. We validated the model through E-cadherin to mimic diffuse type gastric adenocarcinoma, which has a strong predilection for peritoneal spread.

## RESULTS

### Establishment of ex vivo peritoneal metastasis model

Fresh human peritoneal tissue samples were obtained from patients undergoing elective abdominal surgery at Mount Sinai Hospital, Toronto, with REB approval and informed patient consent (Fig. 2A, left panels). On the day prior to surgery (Day −1, Fig. 2B), 10×10^5^ AGS gastric cancer cells that had been dyed red with CellTracker^™^ CM-Dii were seeded into the receded well at the center of a 3.5cm MatTek dish and then cultured overnight in RPMI/10% FBS with Pen/Strep (medium). On the day of surgery, peritoneal tissue was transferred directly from the operative field to a Corning 100 mm Tissue Culture-treated culture dish filled with medium (Fig. 2A, Right panels). Under sterile conditions in the laboratory, peritoneal tissue was divided into 1.5 cm^2^ segments, each of which was placed mesothelial side down onto the MatTek dish over the receded well, without direct contact between the cancer cells and the peritoneal tissue (Fig. 2B, Right). The dish was incubated at 37°C for 24 hours, following which the peritoneal tissue was removed to a fresh 10 cm Petri dish, and washed vigorously on both sides with PBS before being transferred to a new MatTek dish containing medium only, again mesothelial side down. The MatTek dish allows for live cell imaging from below using confocal microscopy. Fluorescence images were obtained with a LEICA SP8 microscope, daily for 5 days (Fig. 2B).

**Figure 2.**
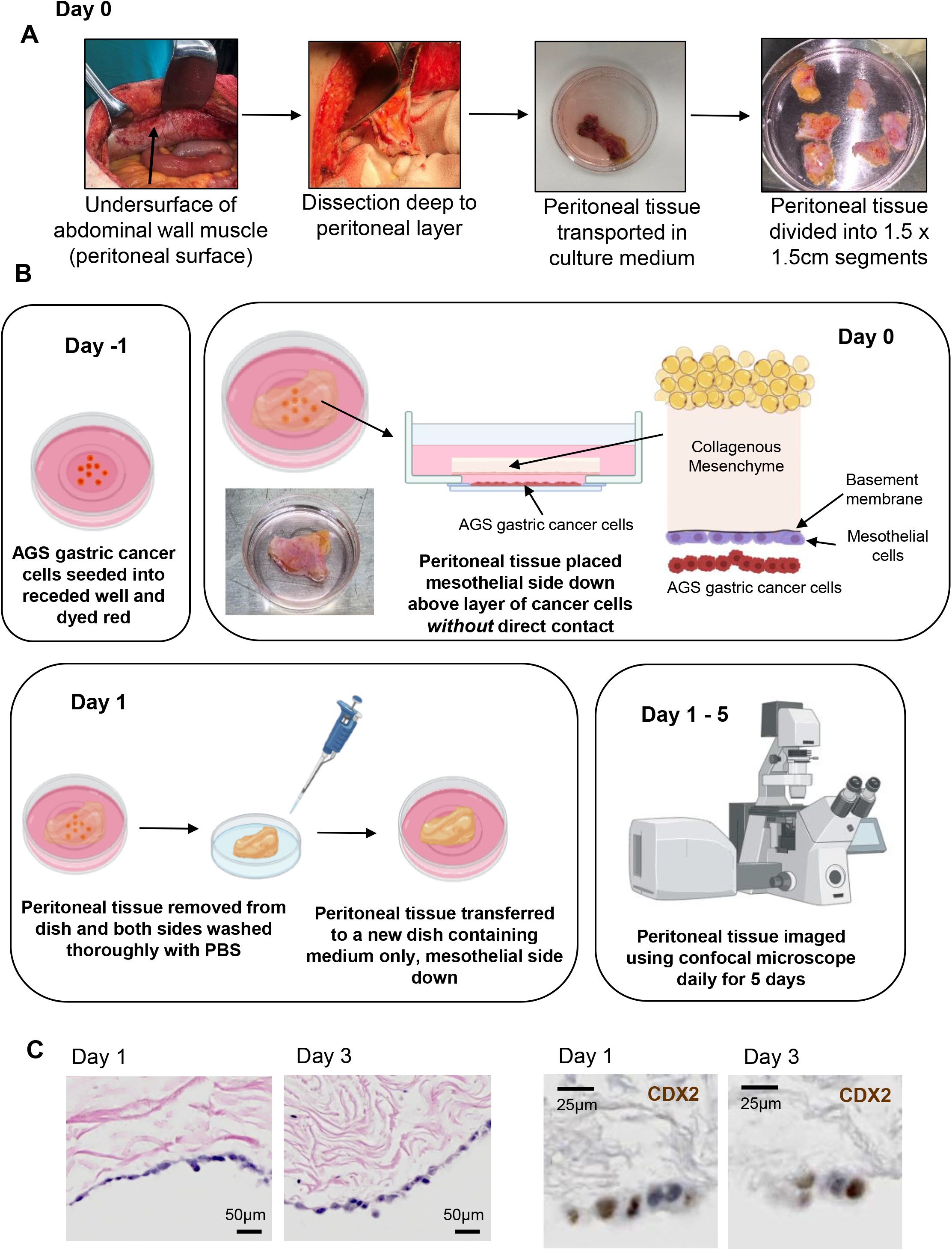
*Ex vivo* human peritoneal metastasis model: Experimental set up and time line. **A)** On day 0, a sample of peritoneal tissue from patient undergoing elective laparotomy in the operating room (OR) and transported to the laboratory in sterile culture medium. **B)** Experimental workflow. Day −1: AGS gastric cancer cells are seeded into receded well of MatTek dish and dyed red using DiI cell tracker (Thermofisher). Day 0: Peritoneal tissue from the OR is divided into segments and each segment placed mesothelial side down over a receded well, with no direct contact between the cells and the tissue. Day 1: Peritoneal tissue is removed from dish and washed thoroughly with PBS before being transferred into a fresh MatTek dish with medium only, mesothelial side down. Images were taken with a confocal microscope daily from Day 1 to 5. In parallel, peritoneal tissue samples were subjected to tissue processing (FFPE) daily (see Supp. Fig. 1A). **C)** Representative images of FFPE sections of *ex vivo* peritoneum at Days 1 and 3, stained with hematoxylin and eosin (H&E) (×20) (left) or stained by immunochemistry for CDX2. AGS cells along the mesothelial surface stain positive for CDX2, a marker for adenocarcinomas derived from an upper gastrointestinal source.

To confirm that the co-cultured explants remained intact and viable over the planned course of the experiment, parallel samples were subjected to conventional histologic examination. Over serial daily intervals, the peritoneal tissue was removed from the MatTek well and processed *in toto* [30]. The entire tissue block was fixed in 10% formaldehyde for 24 hours, then sliced into into thin strips (50mm) at right angles to the peritoneal surface (mesothelial layer, coloured beige in Supp. Fig. 1A). The entire block was cut at 2mm intervals, yielding twenty 5-micron (Supp. Fig. 1B); this ensured that the entire strip of peritoneal tissue was sampled. Stained slides were evaluated by a specialized GI pathologist, blinded to the experiment. Examination of H&E stained sections revealed viable appearing cells of two morphologic types along the mesothelial surface of the explanted peritoneum at days 1 and 3 (Fig. 2C, Left panels). Thin, flat cells were mesothelial cells and plump, round cells with prominent nuclei were AGS cells. This was further confirmed by IHC staining for the gastrointestinal epithelial marker CDX2, which stained AGS cells at days 1 and 3 (Fig. 2C, Right panels).

### Cyto-architecture of explanted human peritoneal tissue is preserved ex vivo

Explanted tissues have a variably limited life span *ex vivo*. Resident cells of the peritoneum inhabit a specific microenvironmental niche that exposes them to peritoneal fluid and that is characterized as relatively hypoxic, hypoglycemic, and hypercarbic [31]. To determine how the *ex vivo* culture conditions used in our model affected the integrity of human peritoneum, we analyzed its structure over the planned experimental time course. Examination of H&E stained sections showed that the cyto-architecture of the peritoneal samples appeared to remain essentially intact over a period of 5 days *ex vivo* (Fig. 3A). The cellular composition of the explanted peritoneum largely mirrored that expected for fresh human peritoneum (Fig, 3B,C) [2]. The submesothelial adipocytes retained their distinct appearance (Fig. 3A), and apparently viable fibroblasts and leukocytes were also observed in the submesothelial layer on Day 3 (Fig. 3C). Intact microvascular and lymphatic structures (Fig. 3D) were observed in the adipocytic layer on Day 3 as well. Delicate mesothelial cells line the peritoneal surface, and are supported on a basement membrane composed of type IV collagen and laminin [1]. An intact layer of mesothelial cells was observed on Days 0 through 2 after peritoneal explantation (Fig. 3E). By Day 3, the mesothelial layer had become more patchy (see staining for calretinin on Day 3, Fig. 3C) and was lost by day 5 (Fig. 3A). This is consistent with the weak attachment of the mesothelial cells to the basement membrane observed *in vivo* [3].

**Figure 3.**
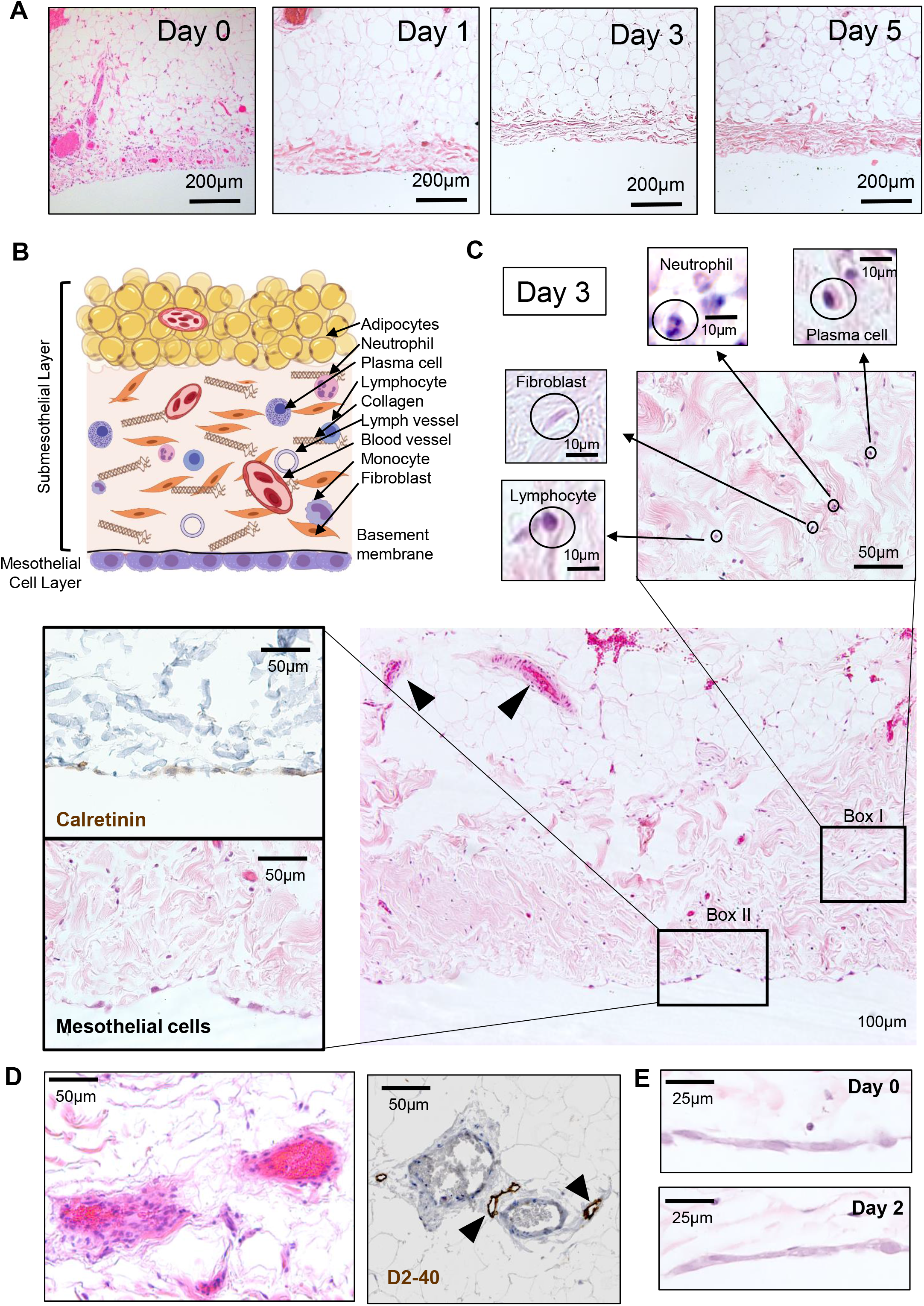
Cyto-architecture of human peritoneum is preserved *ex vivo*. **A)** Representative images of FFPE sections of freshly explanted human peritoneum (Day 0) and on Days 1 through 5 following harvest cultured as described (Fig. 2B) and stained using H&E (x10). Preservation of cyto-architecture over the experimental timeframe is demonstrated. **B)** Cartoon illustrating the cellular and stromal component of normal peritoneum, which is composed of a mesothelial layer (mesothelial cells and underlying basement membrane) and a complex submesothelial layer, which is made up of a largely collagenous mesenchyme and an underlying adipocytic mesenchyme. **C)** On Day 3 post explant, intact blood vessels are observed in the submesothelial layer (arrowheads). Fibroblasts and leukocytes (neutrophils, lymphocytes, plasma cells) are present in the collagenous mesenchyme (Box I). Mesothelial cells stain positive for calretinin (Box II). **D)** Blood vessels in the submesothelial layer are accompanied by lymphatic vessels, which stain positive for D2-40 (arrowheads). **E)** Representative images of FFPE sections of human peritoneal tissue peritoneum stained using H&E (x20) showing intact mesothelial cells at Days 0 (top) and 2 (bottom).

### Assessment of interaction between cancer cells and explanted peritoneum

As illustrated in Fig. 2B, AGS cancer cells were allowed 24h to interact with the explanted peritoneal tissue suspended over them, following which the peritoneal sample was washed thoroughly to remove non-adherent cells. AGS cells that remained “implanted” within the peritoneal tissue were then imaged serially in two separate assays: an Implantation Assay and an Invasion Assay (Fig. 4 A-C). For both assays, the peritoneal tissue was imaged from beneath the mesothelial surface using a confocal microscope to capture a stack of images 2 μm apart, for a total depth of 50-75 μm (z axis). The AGS cancer cells, which had been dyed red, appear distinct against the background green reflectance of the peritoneal tissue’s collagen fibres. For each 1.5 × 1.5 cm peritoneal sample and experimental condition, five different areas of 583 × 583 μm were assessed daily. For the Implantation Assay (Fig. 4A top, Fig. 4B), the entire z stack of images was collapsed into one *en face* image, and the number of AGS cells associated with the peritoneal tissue was measured using FIJI plugin Track Mate [23], by selecting the red channel only. The final Implantation value reported for each day for a given peritoneal sample was the mean of the 5 square areas (Fig. 4B shows an example from Day 2). For the Invasion Assay (Fig. 4A bottom, Fig. 4C), depth of cancer cell invasion into the peritoneal tissue was measured serially over 5 days. The acquired stack of confocal images was reconstructed into a 3D structure that was then analyzed as 512 slices 1.14 μm apart (Y axis). The 512 slices were scanned for the position of the AGS cells relative to the mesothelial surface, and depth of invasion for an individual cell was defined as the distance from the mesothelial surface (shown in fuchsia in Fig. 4C) to the mid-point of the cell (illustrated by tip of white arrows in Fig. 4C) in an orthogonal view. The average depth of three deepest cells for each square area were recorded, and the final Invasion value reported for each day for a given peritoneal sample was the mean of the 5 square areas (Fig. 4C shows an example from Day 5). The latter calculation was done manually by two independent observers and the scores were combined and averaged.

**Figure 4.**
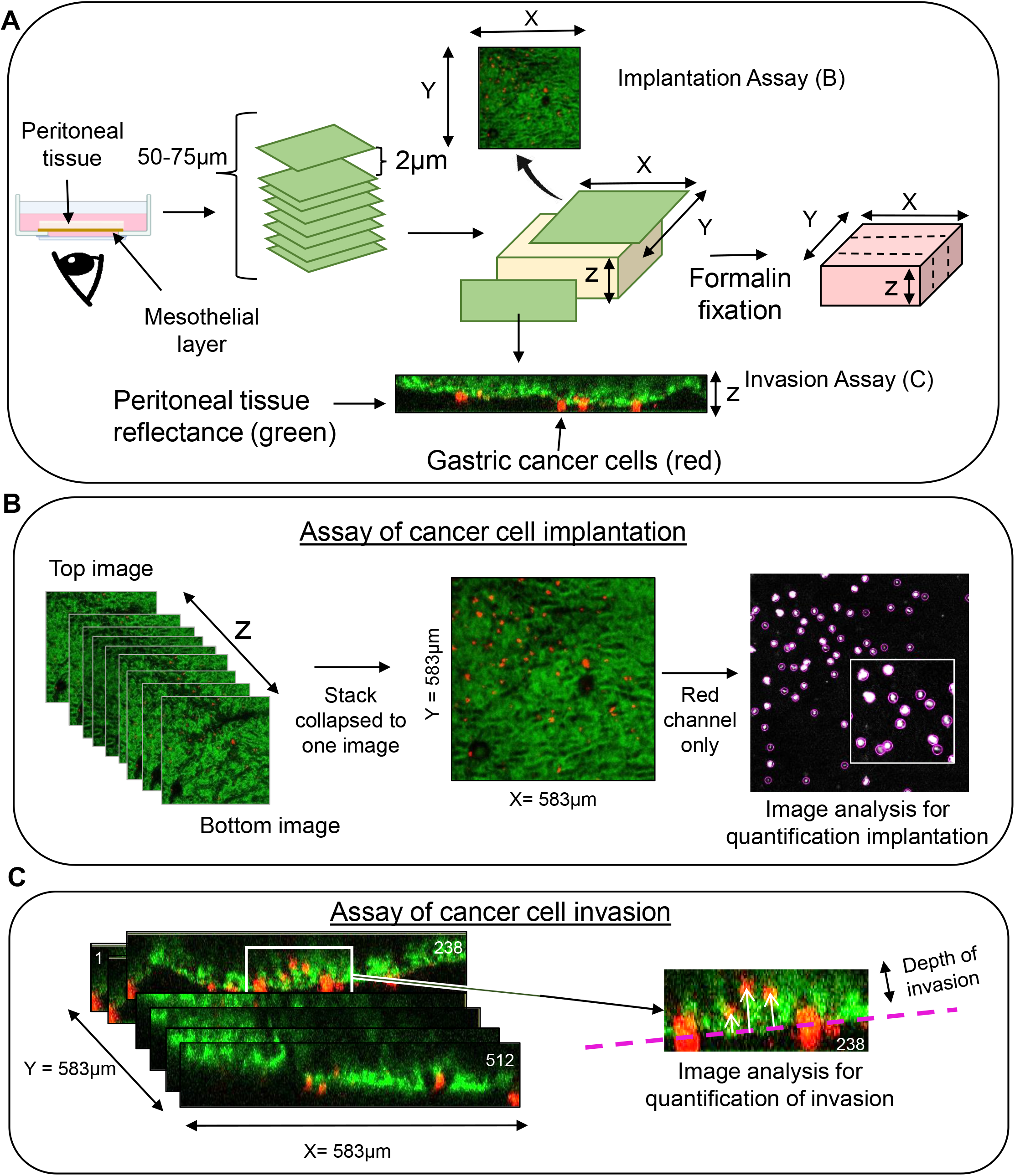
Methods used to assess implantation and invasion of cancer cells into human peritoneal tissue. **A)** Peritoneal tissue was imaged from below the mesothelial surface and a stack of images, 2μm apart, was taken by confocal microscopy (total depth 50-75μm). Background peritoneal tissue appears green (reflectance of collagen fibers), and cancer cells appear red. **B)** To assess cancer cell implantation, the Z-stack of images was collapsed into one image and the number of cells within the peritoneal tissue was measured by selecting the red channel only. The number of cells was determined using FIJI plugin TrackMate (see methods). **C)** To assess depth of cancer cell invasion into peritoneal tissue, distance between the mesothelial cell layer (fuchsia dashed line) and the mid-point of the cell was measured on orthogonal view (FIJI) and the mean of the 3 deepest cells per field was calculated (see methods).

### Ex vivo model is sensitive to functional effects of gene manipulation

To determine whether the explant model would permit detection of alterations in peritoneal implantation and invasion by AGS cells, we reprogrammed the latter to restore an epithelial phenotype. Parental AGS cells do not express functional E-Cadherin (CDH1) due to a truncation mutation (p.T578fs) [32]. Using lentiviral infection of AGS cells, we stably restored wild-type CDH1 (Fig. 5A), and observed the expected reversal of the more mesenchymal phenotype seen in V5-alone controls (Fig. 5B). Restoration of function CDH1 also suppressed 2D migration (Fig. 5C), without altering AGS cell viability or proliferation (Supp. Fig. 2A, B). In co-culture with fresh human peritoneal tissue, CDH1 restoration dramatically suppressed both implantation (Fig. 6A) and invasion (Fig. 6B) into the explanted peritoneum. This result confirmed the sensitivity of the model to manipulation of cancer cell genotype/phenotype, demonstrating its viability in studying the molecular determinants of peritoneal metastasis. The sequential increase in the number of V5-only expressing cells observed within the peritoneum between days 1 and 3 suggests that the implanted AGS cells were proliferating.

**Figure 5.**
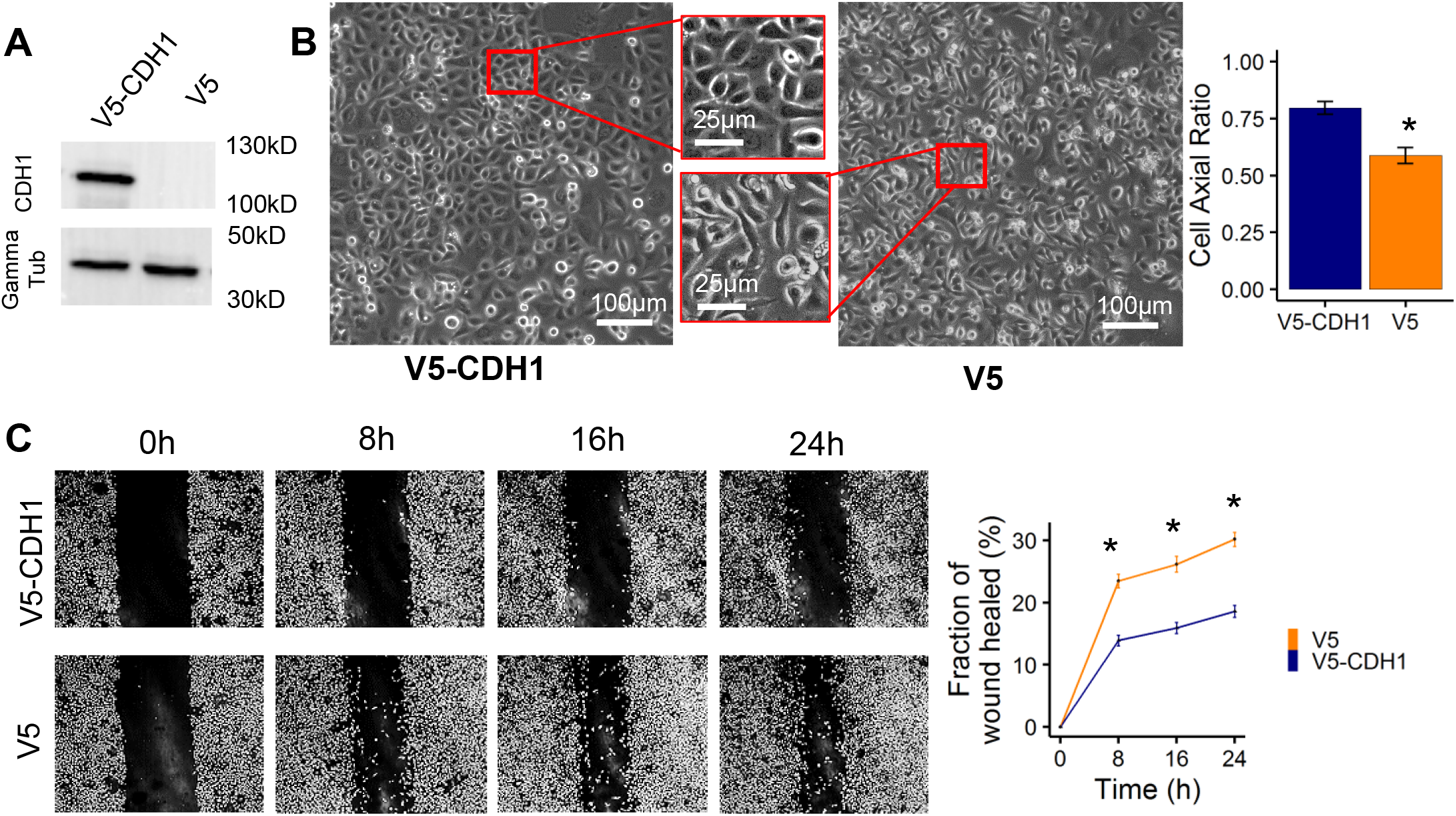
Restoration of functional E-cadherin (CDH1) increases cell-cell adhesion and suppresses migration in AGS gastric cancer cells. **A)** Extracts from AGS cells expressing V5-tagged CDH1 or V5 alone were subjected to immunoblot, using anti-CDH1 antibody. **B)** Brightfield imaging showing that restoration of functional CDH1 in AGS cells caused aggregation due to increased cell-cell adhesion (left). Quantification of cell shape where 1 represents a perfect circle: shows that cell roundness is increased in AGS cells expressing functional CDH1 versus V5 alone (right). **D)** Directional migration, assessed in a wound healing assay, is reduced in AGS cells expressing functional CDH1 versus V5 alone. Summary of 3 independent experiments (right). Data are presented as mean ± SEM of 3 independent experiments, *=p<0.001.

**Figure 6.**
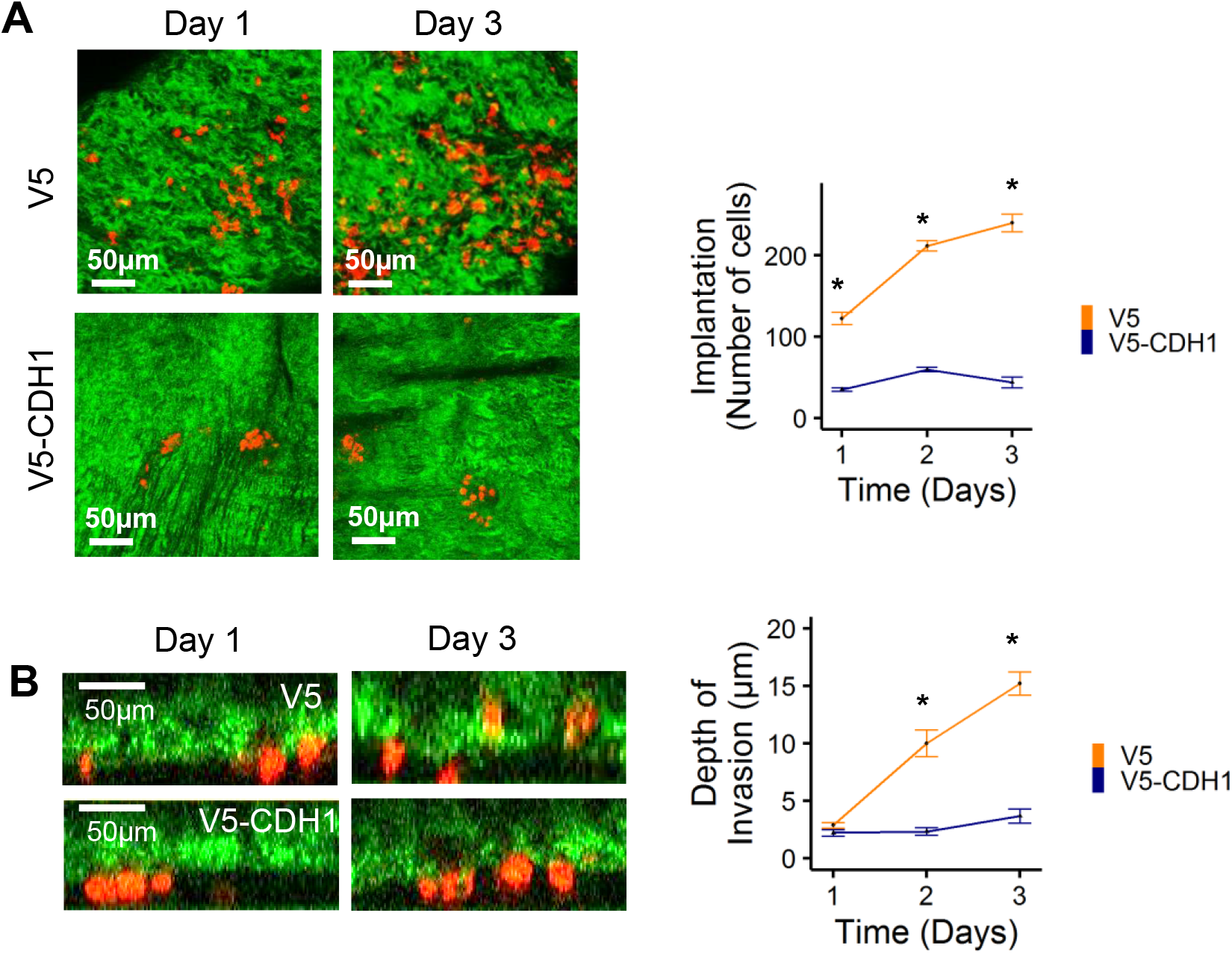
Restoration of functional E-cadherin (CDH1) suppresses implantation and invasion into the peritoneum by AGS cancer cells. **A)** En-face fluorescence time-lapse images (left) of *ex vivo* peritoneal tissue showing presence of AGS cells expressing V5-CDH1 or V5 alone within the peritoneum (implantation). Quantification of number of cells per high power field (HPF, 583μm x 583μm) over time shows a marked reduction in implantation in CDH1 expressing cells (right). Data are presented as mean ± SEM of 3 independent experiments, *p<0.001. **B)** Cross sectional fluorescence time-lapse images (left) of *ex vivo* peritoneal tissue showing depth of invasion by AGS cells expressing V5-CDH1 or V5 alone. Quantification of depth of invasion over time shows reduced impaired invasion with V5-CDH1 expression. Data are presented as mean ± SEM of 3 independent experiments, *p<0.001.

### Human cell lines screened for implantation in peritoneal explant mode.l

We tested a variety of human cell lines for peritoneal implantation (Supp. Fig. 3A, B). The non-malignant line RPE did not implant, and neither did the soft tissue sarcoma line Ht1080 or the osteosarcoma line U2OS. Carcinoma lines derived from lung (Adenocarcinoma AC549), oropharynx (Squamous cell carcinoma UMSCC1) and urinary bladder (Transitional cell carcinoma T24) also failed to implant. By contrast, cell lines derived from cancer types that are known to metastasize to the peritoneal cavity in patients, including adenocarcinoma of the ovary (HEY), cervix (Hela), pancreas (Panc1) and colorectal (Hct116) did implant within the explanted human peritoneum. Thus, the model could discriminate between cancer types that are prone to peritoneal dissemination and those that are not.

## DISCUSSION

Here, we have established an *ex vivo* co-culture system that uses freshly harvested human peritoneum as a tool to investigate implantation of gastric cancer cells through the peritoneal mesothelium and invasion into the submesothelial space. This system models steps of the peritoneal metastatic cascade. This model overcomes several limitations of *in vitro* and *in vivo* model systems. The cytoarchitecture of the explanted peritoneal tissue is retained, both recapitulating the physical and cellular barriers to implantation and invasion present *in vitro*. The model allows investigation of gene expression profiles that enable or suppress peritoneal metastasis, as demonstrated by restoration of wild type CDH1 gene product in AGS cells with resultant disabling of peritoneal implantation.

As far as we are aware, only one other *ex vivo* human peritoneal model that investigates gastric cancer dissemination has been published to date. In that model, which used peritoneum taken with a hernia sac during elective hernia repair, gastric cancer cells were seeded densely onto the peritoneum [33]. In the model we describe here, gastric cancer cells are not in physical contact with the peritoneum, since they are cultured in a monolayer at the bottom of a receded well, at a distance of 750 microns from the peritoneal mesothelium. This platform requires active movement of cells within a liquid microenvironment and against the forces of gravity, mimicking *in vivo* forces that exist. Other distinguishing features of our model are the histologic verification of maintenance of normal peritoneal cytoarchitecture, and real time serial fluorescence imaging of the cancer cells as they move into and within the peritoneal surface. The cell tracker dye becomes membrane impermeant once it is within the cell, and continue to fluoresce over 3-6 subsequent cell divisions, meaning that the cell and its progeny can be followed over time [34]. The peritoneal tissue is visualized through its endogenous reflectance, meaning no fixation or staining is required [35]. The automated quantification of cellular implantation using FIJI trackmate eliminated observer bias and inter-observer variability [36].

Having established the sensitivity and discriminatory capabilities of the *ex vivo* model described here, we anticipate various extension of the experimental design that would enable more detailed investigation of several aspects of the peritoneal metastatic cascade. Initial attachment/ adhesion to the peritoneal mesothelium and interaction with the mesothelial cells could be studied within the first few hours of co-culture. Cancer cells that implant could be compared to those that do not to screen for enabling expression profiles in an unbiased manner. Drugs and other compounds with the potential to target specific aspects of the peritoneal metastatic could be screened [37], and deleterious side effects on the peritoneal tissue itself could potentially be detected.

We had anticipated challenges with reproducibility of results, given that the peritoneal tissue samples we used here came from a wide variety of patients who were undergoing abdominal surgery. Their median age was 61, but ranged from 28 to 80 (n =8), female to male ratio was 5:3. In no case was there any macroscopic abnormality of the sampled peritoneum. While there may have been inter-patient variability in terms of submesothelial stroma thickness and composition [3], the depth of penetration of the cancer cells over the time line studied was rather limited, and this may have favoured reproducibility of results. This point emphasizes the applicability of this model to the early phases of peritoneal metastasis, before the implanted cells proliferate into a colony, acquire a blood supply and engender a fibrotic response, as seen in the well-established peritoneal metastasis shown in Fig. 2B [38]. The main advantage of the explant model is the ability to focus on these initial processes and what enables them. In terms of implications for discovery science and clinical translation, the *ex vivo* peritoneal metastasis model we describe here offers the capacity to screen genes/ pathways of interest, as well as novel therapies in a relatively simple co-culture system that recapitulates key aspects of the *in vivo* microenvironment. It is also applicable to other cancer types that harbour a predisposition to peritoneal dissemination, such as ovarian, pancreatic and colon cancer [5, 39].

## METHODS

### Cell culture

Cells were grown at 37°C in DMEM (HeLa, Hct116, HEY, T24), RPMI1640 (Ht1080, Panc1, RPE), F12K (AGS) or McCoy 5A medium (USMCC1, U2OS) supplemented with 10% FBS (Wisent). AGS, USMCC1, HEY, Ht1080, and HeLa cell lines were obtained from ATCC in 2018, 2016, 2013, 2010 and 2009, respectively, and were maintained between passages 4 and 15. The U2OS line was a kind gift from the Laurence Pelletier laboratory (obtained in 2013 and maintained between passages 6 and 10; Lunenfeld Tanenbaum Research Institute, Sinai Health System, Toronto, Canada). Panc1, Hct116, T24 and RPE lines were a kind gift from the Daniel Schramek laboratory (obtained in 2020 and maintained between passages 8 and 28; Lunenfeld Tanenbaum Research Institute, Sinai Health System, Toronto, Canada). Cell lines were not further authenticated in our laboratory.

### Stable cell lines

AGS cells, which are derived from primary gastric adenocarcinoma, do not express CDH1 protein due to a truncating mutation (T578fs) [32]. The parental AGS cell line was infected with lentivirus to stably express V5-CDH1, or V5 alone, using gateway constructs (pLEX_306m, Addgene). Lentiviruses were produced as described [40] and used to infect the cells for 24h, followed by puromycin (2ug/ml) selection for 7 days. To confirm protein expression, cell extracts were separated by SDS-PAGE, transferred onto PVDF membranes, blocked with 5% milk-PBS-0.1%Tween, probed with CDH1 (BD Transduction Laboratories, 610182, 1:1000) at 4°C overnight, then HRP-linked secondary antibody (GE Healthcare), and detected using SuperSignal West Femto Maximum Sensitivity Substrate (34095; Thermo Fisher Scientific).

### Measurement of cell viability

30 × 10^5^ V5 or V5-CDH1 expressing AGS cells were seeded on a clear flat bottom 96-well black polysterene plate (Corning). Viability was assessed using the LIVE/DEAD^®^ Viability/Cytotoxicity Kit (ThermoFisher), according to the manufacturer’s instructions. Cells were viewed using the INCell6000 analyzer equipped with a live cell apparatus (37°C humidified chamber with 5% CO2) with a 96-well plate adaptor. Automated data acquisition was at the indicated time points. Images were collected with a 10X objective lens. All hardware and image capture conditions were enabled and images analyzed by MATLAB 9.4 2018.

### Assessment of cell morphology

Images of cells grown to ^~^80% confluence on the 96-well plates were captured by INCell Analyzer 2000. FIJI was used to analyze the images and determine cell axial ratio (width/length). 30 × 10^5^ V5 or V5-CDH1 expressing AGS cells were imaged per well, 20 fields per well.

### Cell staining for live cell imaging

AGS cells plated on MatTek dishes were stained using CellTracker^™^ CM-DiI (ThermoFisher, C7000). AGS cells were first plated on MatTek dishes for 18h prior to staining. Media was removed from cells and cells were incubated with dye (1 μg/μl) for 30 minutes.

### Histologic and Immunohistochemical (IHC) evaluation

Tissue samples were fixed in 10% formalin for 24h at room temperature. Serial coronal sectioning was performed at three levels, 50 mm apart. Each tissue edge was stained red with Tissue-Marking dye to orientate the specimen for paraffin embedding (Supp. Fig. 1A). The strips of peritoneal tissue were paraffin embedded by lining the three strips parallel to each other, with the mesothelial layer oriented to one side (Supp. Fig. 1B). Sections (5μm) were stained with hematoxylin and eosin (H&E) according to standard protocols using the Veristain Gemini Automated Slide Stainer (Thermo Scientific) at the University of Toronto Dentistry Pathology Core, and examined with a Leica DMR upright microscope. For IHC staining, slides were stained with CDX2 or D2-40 antibodies using the fully automated Dako Omnis platform (deparaffinization and retrieval built in), using the Dako OMNIS detection kit (Ref: GV800). Slides were exposed to high pH for 15 minutes, antibody incubation for 10 minutes, polymer detection for 20 minutes, substrate chromogen for 5 minutes and finally Dako hematoxylin for 3 minutes. This was performed at the Department of Pathology at the Hospital for Sick Children, Toronto. An experienced pathologist rated staining as strongly or weakly positive, or negative, on a per cell basis.

### Scratch wound migration

1×10^6^ cells were seeded into 6 well plates and serum starved for 18h prior to scratch. Scratches were made manually using a P10 pipette tip, and migration was assayed for 24h using the INCell6000 analyzer equipped with a motorized stage and a live cell apparatus (37°C heated and humidified chamber with 5% CO2) with a 6-well plate adaptor. Data acquisition was performed continuously over the indicated time courses. Images were collected with a 10x objective lens. All hardware and image capture conditions were made possible, and images analyzed, using MATLAB 9.4 2018.

Image analysis was carried out by measuring the total wound area in three fields per condition, at t=0, 8, 16 and 24 h. Measurements at each location were averaged to yield a mean wound area. The residual wound area was expressed as a percent of original wound, which was itself highly reproducible, subtracted from 100 to yield the %healed at each time point, and compared across groups by t-test. The Bonferroni correction sets the significance at P<0.05 (two tailed).

