## Supplemental Figures 1-3 for "A Novel *Ex Vivo* Peritoneal Model to Investigate Mechanisms of Peritoneal Metastasis in Gastric Adenocarcinoma"

### Supplemental Figure 1

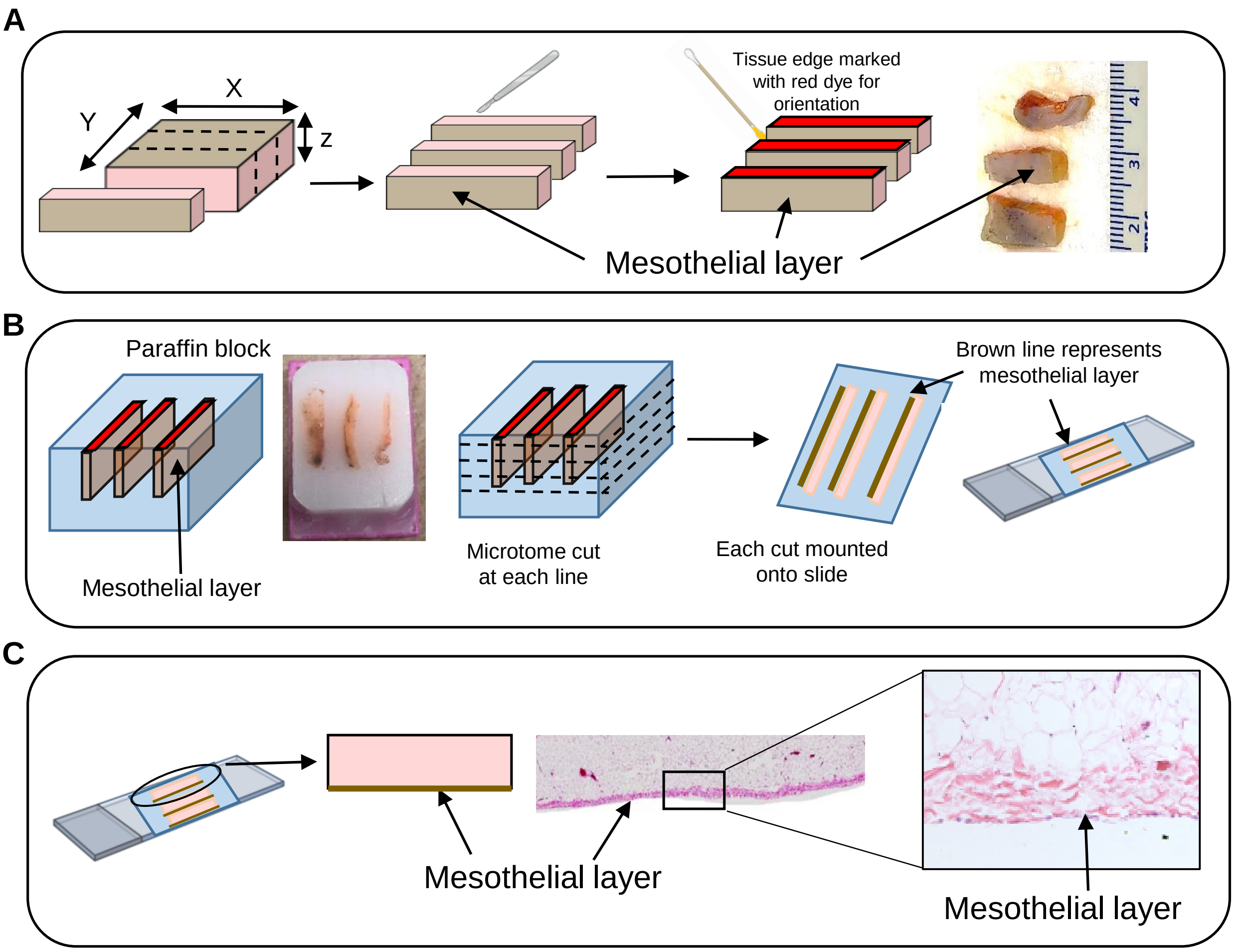

Supplemental Figure 2

**A**

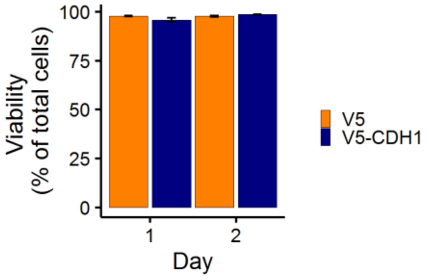


**B**

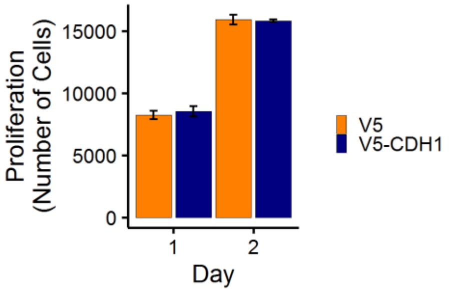

### Supplemental Figure 3

**A**

**Cells lines that implanted**

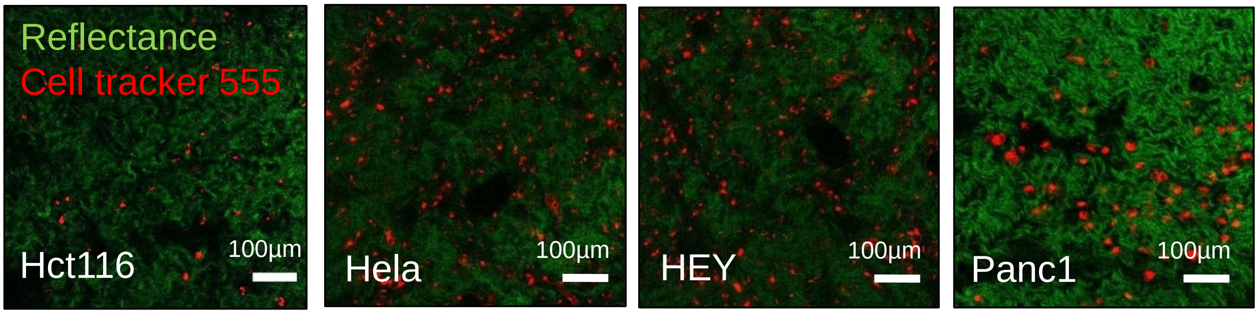


**B**

**Cells lines that did not implant**

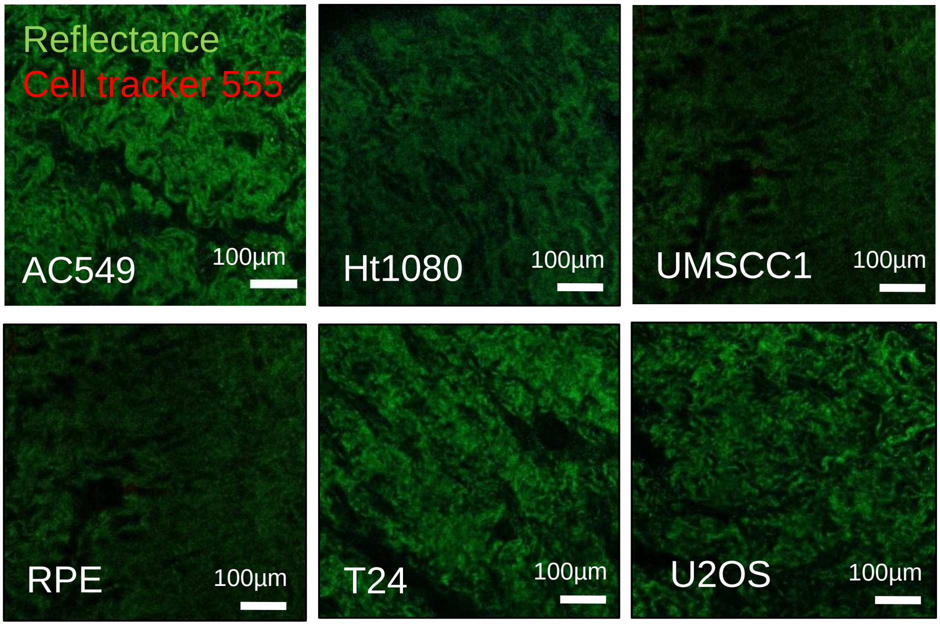

#### SUPPLEMENTAL FIGURE LEGENDS

**Supplemental Figure 1. Methods used for sectioning peritoneal tissue. A)** The entire peritoneal tissue sample was sliced into blocks (1/3 of Y x Z x X, 5 x 5 x ~30mm) at right angles to the mesothelial layer (beige), and stained with red dye for orientation in preparation for paraffin embedding. **B)** The three thinner strips of peritoneal tissue were cute, then paraffin embedded in parallel, mesothelial layer facing one side. Twenty 5µm cuts were then obtained from each paraffin block, spanning the entire paraffin block, thereby ensuring that the entire strip of peritoneal tissue was sampled. Each 5µm thick cross section was mounted onto a glass slide and inspected. **C)** H&E staining of the sections of peritoneal tissue, demonstrating orientation of mesothelial surface.

**Supplemental Figure 2. Restoration of functional E-cadherin (CDH1) did not affect viability or proliferation of AGS cancer cells. A)** >95% of AGS cells remained viable over 2 days in culture, with no difference between V5-CDH1and V5 alone controls. Data are presented as mean ± SEM of 3 independent experiments. **B)** Proliferation of AGS cells under culture conditions used in explant model was equivalent in V5-CDH1 and V5 AGS cells, as measured by number of cells per day. Data are presented as mean ± SEM of 3 independent experiments.

**Supplemental Figure 3. Human cell lines screened for implantation in peritoneal explant model. A)** Implantation of HCT116 (colorectal cancer), HeLa (cervical cancer), HEY (ovarian cancer) and PANC-1(pancreatic cancer) into explanted human peritoneum. Representative images taken by confocal microscopy on day 2 of the experiment. **B)** There was no apparent implantation of AC549 (Lung cancer), HT1080 (fibrosarcoma), UMSCC1 (oral cancer), RPE (retinal pigment epithelial cells), T24 (bladder cancer) or U2OS (osteosarcoma) into explanted human peritoneum. Representative images taken by confocal microscopy on day 2 of the experiment.
